# Combining repetition suppression and pattern analysis provides new insights into the role of M1 and parietal areas in skilled sequential actions

**DOI:** 10.1101/2020.08.21.261453

**Authors:** Eva Berlot, Nicola J. Popp, Scott T. Grafton, Jörn Diedrichsen

**Affiliations:** The Brain and Mind Institute, University of Western Ontario, London, ON N6A 5B7, Canada; Department of Psychological and Brain Sciences, University of California, Santa Barbara, CA 93106, USA; Institute for Collaborative Biotechnologies, University of California, Santa Barbara, CA 93106, USA; Department of Statistical and Actuarial Sciences, University of Western Ontario, London, ON N6A 5B7, Canada; Department of Computer Science, University of Western Ontario, London, ON N6A 5B7, Canada

## Abstract

How does the brain change during learning? In functional magnetic resonance imaging studies, both multivariate pattern analysis and repetition suppression (RS) have been used to detect changes in neuronal representations. In the context of motor sequence learning, the two techniques have provided discrepant findings: pattern analysis showed that only premotor and parietal regions, but not primary motor cortex (M1), develop a representation of trained sequences. In contrast, RS suggested trained sequence representations in all these regions. Here we applied both analysis techniques to a 5-week finger sequence training study, in which participants executed each sequence twice before switching to a different sequence. Both RS and pattern analysis indicated learning-related changes for parietal areas, but only RS showed a difference between trained and untrained sequences in M1. A more fine-grained analysis, however, revealed that the RS effect in M1 reflects a fundamentally different process than in parietal areas. On the first execution, M1 represents especially the first finger of each sequence, likely reflecting preparatory processes. This effect dramatically reduces during the second execution. In contrast, parietal areas represent the identity of a sequence, and this representation stays relatively stable on the second execution. These results suggest that the RS effect does not reflect a trained sequence representation in M1, but rather a preparatory signal for movement initiation. More generally, our study demonstrates that across regions RS can reflect different representational changes in the neuronal population code, emphasizing the importance of combining pattern analysis and RS techniques.

**Significance statement:** Previous studies using pattern analysis have suggested that primary motor cortex (M1) does not represent learnt sequential actions. However, a study using repetition suppression (RS) has reported M1 changes during motor sequence learning. Combining both techniques, we first replicate the discrepancy between them – with learning-related changes in M1 in RS, but not pattern dissimilarities. We further analysed the representational changes with repetition, and found that the RS effects differ across regions. M1’s activity represents the starting finger of the sequence, an effect that vanishes with repetition. In contrast, activity patterns in parietal areas exhibit sequence dependency, which persists with repetition. These results demonstrate the importance of combining RS and pattern analysis to understand the function of brain regions.

## Introduction

The ability to learn and produce complex sequences of movements is essential for many everyday activities, from tying shoelaces to playing instruments. Searching for where these acquired skills are represented in the brain has been one of the central questions in motor neuroscience (Lashley, 1950). One prominent issue in this debate is whether skilled sequence execution relies on representations in premotor and supplementary motor areas, or whether the sequences are represented in the primary motor cortex (M1) (see Dayan and Cohen, 2011; Berlot et al., 2018 for reviews). We recently conducted a systematic longitudinal 5-week training study (Berlot et al., 2020) employing functional magnetic resonance imaging (fMRI) to assess brain changes with motor sequence learning. We observed no overall change in overall activity with learning in M1, and no changes in the sequence-specific activity patterns. In contrast, clear learning-related changes in both overall activity and fine-grained activity patterns were observed in premotor and parietal areas, suggesting learning-related changes occur outside of M1. Consistent with this idea, activity patterns in M1 seem to reflect individual movement elements, but not the sequential context (Yokoi et al., 2018; Yokoi and Diedrichsen, 2019; Russo et al., 2020). This suggests that M1 does not represent learnt motor sequences, but must rely on inputs from other areas to select the next correct movement element.

Using the technique of repetition suppression, however, Wymbs and Grafton (2015) provided evidence for learning-related changes during motor sequence learning in M1. Repetition suppression (RS) refers to the observation that a stimulus repetition evokes reduced neuronal activity compared to its initial presentation (Gross, Schiller, Wells, Gerstein, 1967). It is commonly used as a tool for investigating brain representation (Buckner et al., 1998; Henson et al., 2003; see Segaert et al., 2013 for review) following the logic that if regional activation reduces upon repetition, the underlying neuronal population must represent some aspect of the stimulus that repeated (Grill-Spector et al., 2006). Wymbs and Grafton (2015) found learning-related changes in RS across several regions, including M1, where they reported a non-monotonic change in RS over weeks – early increase, followed by a decrease, and again an increase in RS, which they suggested indicates skill-specific specialization in M1. Altogether, their results indicate that M1’s activity patterns are malleable when learning motor sequences. This stands in stark contrast to the above-mentioned studies that used pattern dissimilarity analyses and found no evidence of sequential representation in M1.

We reasoned that this discrepancy between RS and pattern analysis may reflect the fact that different underlying components of activity patterns might bring about the suppression of activity observed on repetition, some of which may not be directly related to a sequence identity (Grill-Spector et al., 2006; Alink et al., 2018). To understand RS effects in more detail, we need to know what aspects of the underlying representations reduce from the first to the second repetition. We therefore designed a paradigm that allowed us to investigate changes in brain representation using both tools – RS and multivariate pattern analysis. We trained healthy volunteers to produce motor sequences over 5 weeks and tested their performance during high-field (7 T) MRI scanning. Participants performed trained and untrained sequences, each sequence twice in a row, allowing us to conduct both pattern and RS analysis on the same data. Replicating previous results, we observed significant learning-related changes in M1 for RS, but not for pattern dissimilarities. In contrast, both metrics showed learning-related changes in premotor and parietal regions. Using pattern analysis, we then decomposed the activation patterns in the first and second repetition to determine which representational aspects underlie the RS effects in the different regions. Finally, we performed control analyses to test whether observed effects could be attributed to learning-related improvements in the execution speed.

## Materials and Methods

### Participants

Twenty-seven participants took part in the experiment. Data of one participant were excluded because the field map was distorted in one of the four scans, resulting in 26 participants whose data was analyzed (17 females, 9 males). Their mean age was 22.2 years (SD = 3.3 years). Criteria for study inclusion were right-handedness and no prior history of psychiatric or neurological disorders. They provided written informed consent to all procedures and data usage before the study started. The experimental procedures were approved by the Ethics Committee at Western University.

### Apparatus

Finger sequences were performed using a right-hand MRI-compatible keyboard device (**Fig 1a**). The keys of the device had a groove for each fingertip, with keys numbered 1-5 for thumb-little finger. The keys were not depressible, so participants performed isometric finger presses. The force of the presses was measured by the force transducers underneath each finger groove (FSG-15N1A, Sensing and Control, Honeywell; dynamic range 0-25 N; update rate 2 ms; sampling 200 Hz). For the key to be recognized as pressed, the applied force had to exceed 1 N.

**Figure 1.**
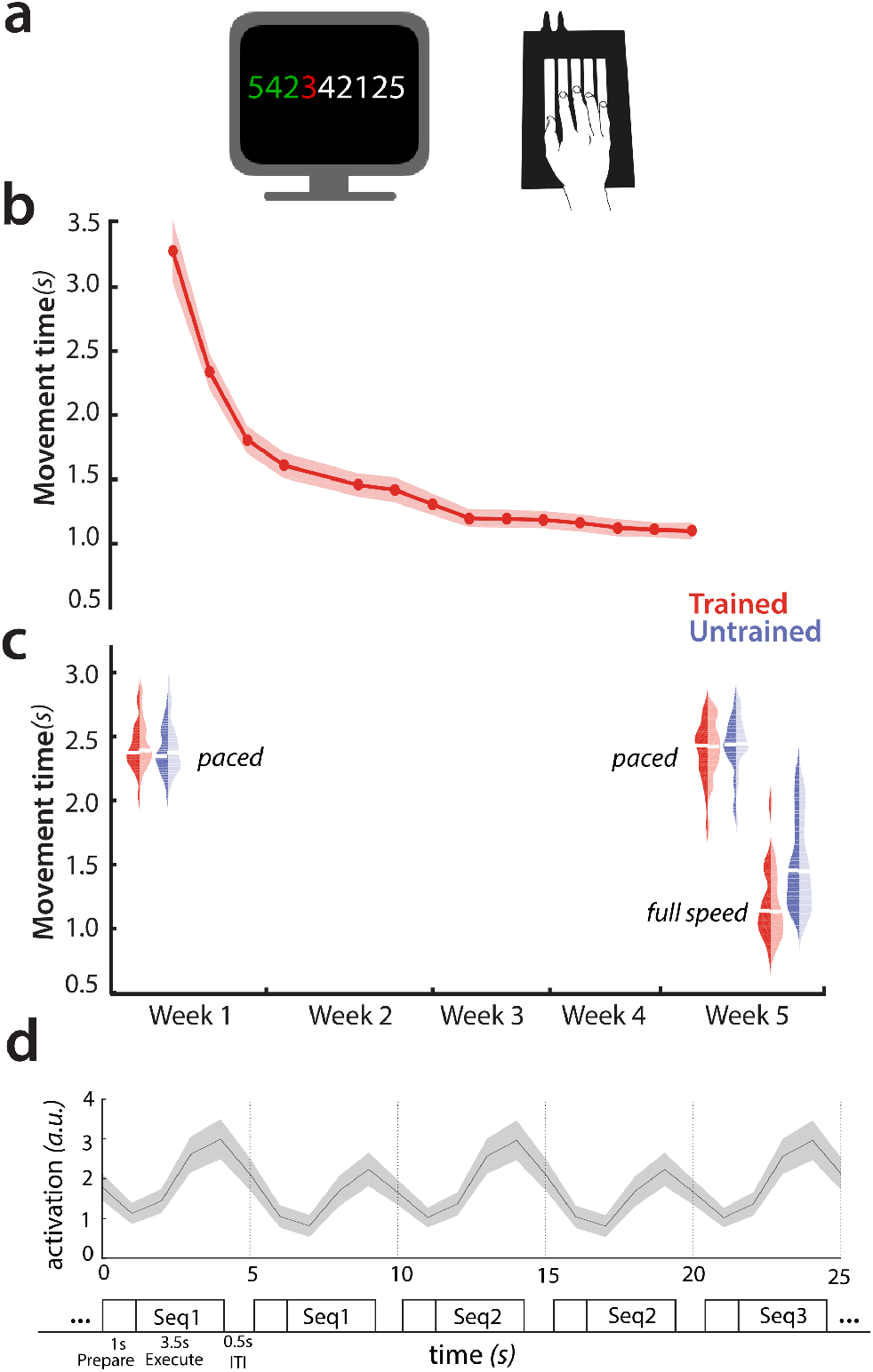
Experimental paradigm. **a)** Experimental setup – finger sequences composed of 9 digits were executed on a keyboard device. Participants received visual feedback on correctness of their presses – digits turned green for correct presses, red for incorrect presses. **b)** Group-averaged performance on trained sequences over the 5-week behavioural training protocol. Red shade indicates the standard error of the group mean. **c)** Group-averaged performance during the scanning sessions. Trained sequences are in red, untrained in blue. Dark colour indicates first execution, light second execution. White bars indicate the group mean performance. **d)** Experimental paradigm inside the scanner. Each sequence was presented twice in a row. Trials started with a 1-s preparation time in which the sequence was presented, followed by a 3.5s-period of main phase, when the sequence was also execution, followed by 0.5 s of inter-trial interval (ITI). The plotted timeseries for an insert of the design is group-averaged evoked activation of M1. Shaded error bars indicate the standard error of the mean.

### Experimental design – learning paradigm

Participants were trained over a five-week time period to perform six 9-digit finger sequences (**Fig 1b**). They were split into two groups, with trained sequences of one group being the untrained sequences of the second group, and vice versa (see **Fig 4b** for all of the chosen sequences). The chosen sequences for both groups were matched as closely as possible on several features: starting finger, number of repetitions per finger, and first-order finger transitions. The decision to split participants into two groups was made to ensure that none of the observed effects could be due to the specific set of sequences chosen.

On day 1 of the study, participants were acquainted with the apparatus and the task performed in the scanner. To ensure no sequence-specific learning would take place prior to scan 1, we used finger sequences different from the trained and untrained sets which participants did not encounter at any later stage of the experiment.

During the behavioral training sessions, participants were trained to perform the six sequences. They received visual feedback on the correctness of their presses online with each digit turning green for correct, and red for incorrect press (**Fig 1a**). They were instructed to perform the sequences as fast as possible while keeping the overall accuracy >85%. The details of the training protocol, as well as a few other design features (which were not assessed for this paper) have been described elsewhere (Berlot et al., 2020).

### Experimental design – scanning

Longitudinal studies assessing learning have to tackle the challenge that performance changes with learning, and that it is not clear whether brain changes reflect the acquisition of new skills, or are caused indirectly by the changed behaviour (Poldrack, 2000). For motor learning, the higher speed of execution could lead to different brain activation, unrelated to learning. Pacing participants to perform at the same speed for trained and untrained sequences, and across sessions, presents a possible solution for this problem. On the other side, pacing participants at a slower speed might not tap into the same neural circuitry as skilled behaviour. For this reason, we decided to include both approaches; sessions with paced performance and a session where participants performed at full speed.

Participants underwent a total of 4 MRI scanning sessions (**Fig 1c**) while executing trained and untrained sequences. The first session served as a baseline prior to the start of the training protocol (in week 1), where the “trained” and “untrained” sequences were both untrained and seen for equivalent amounts of time. The second session was conducted in week 2, and the last two after training protocol was completed – in week 5. In scanning sessions 1-3, participants’ performance inside the scanner was paced with a metronome, whereas in session 4, they performed as quickly as possible. For the purpose of this paper, we analyzed data of scanning session 1 (prior to training – paced), 3 (after learning – paced) and 4 (after learning – unpaced) (**Fig 1c**), allowing us to examining learning- and performance-related changes. Session 4 allows for the closest comparison to the previous RS study (Wymbs and Grafton, 2015) which also employed a full-speed performance design.

Each scanning session consisted of eight functional runs with event-related design randomly intermixing trials containing the 6 trained and the 6 untrained sequences (totalling 72 trials per functional run). Each sequence was executed for two trials in a row (**Fig 1d**). In this way, our design did not differentiate between repetition suppression and expectation suppression (Summerfield et al., 2008; Kok et al., 2012). In contrast to perceptual studies, however, in motor studies the influence of the expectation of a repetition is likely much less important. After the informative cue, preparatory processes are executed in a full awareness of whether the sequence is repeated from last trial, no matter if that repetition was expected or not. Thus, repetition effects in motor control will always contain an element of expectation. For this reason, we chose repetition to be a predictable feature of our experimental design.

Each trial started with a 1-s preparation time with nine digits of the sequence presented on the screen (**Fig 1d**). A ‘go’ signal was presented afterwards. In scans 1-3, a pink line appeared underneath the sequence and started expanding, indicating the pace at which participants were to press. In scan 4, participants executed the sequence as fast as possible after the go cue. After execution, they received feedback on their overall performance – 3 points for correct and 0 for incorrect performance. Each trial lasted for 5 s total, with a 0.5-s inter-trial interval (**Fig 1d**). Five periods of 10 s rests were added throughout each functional run to provide a better estimate of baseline activation. These rests were added randomly, but never between the first and second execution of the same sequence. In total, each scanning session lasted for approximately 75 minutes.

### Image acquisition

Data were acquired on a 7-Tesla Siemens Magnetom MRI scanner with a 32-receive channel head coil (8-channel parallel transmit). At the beginning of the first scan, we acquired anatomical T1-weighted scan for each participant. This was obtained using a magnetization-prepared rapid gradient echo sequence (MPRAGE) with voxel size of 0.75×0.75×0.75 mm isotropic (field of view = 208 x 157 x 110 mm [A-P; R-L; F-H], encoding direction coronal). Data during functional runs were acquired using the following sequence parameters: GRAPPA 3, multi-band acceleration factor 2, repetition time [TR] = 1.0 s, echo time [TE] = 20 ms, flip angle [FA] = 30 deg, slice number: 44, voxel size: 2×2×2 mm isotropic. To estimate magnetic field inhomogeneities, we acquired a gradient echo field map with the following parameters: transversal orientation, field of view: 210 x 210 x 160 mm, 64 slices, 2.5 mm thickness, TR = 475 ms, TE = 4.08 ms, FA = 35 deg. The dataset is publicly available on OpenNeuro (accession number ds002776).

### Preprocessing and first level analysis

Data preprocessing was carried out using SPM12. Preprocessing of functional data included correcting for geometric distortions using the acquired field map data, and head motion correction (3 translations: x, y, z; 3 rotations: pitch, roll yaw). The data across sessions were all aligned to the first run of the first session, and then co-registered to the anatomical scan.

Preprocessed data were analysed using a general linear model (GLM; Friston et al., 1994). We defined a regressor for each of the performed 12 sequences (6 trained, 6 untrained), separately for their first and second execution – resulting in a total of 24 regressors per run. The regressor was a boxcar function defined for each trial, and convolved with a two-gamma canonical hemodynamic response function (time to peak: 5.5 s, time to undershoot: 12.5 s). All instances of sequence execution were included into estimating regressors, regardless of whether the execution was correct or erroneous. This analysis choice was also taken by Wymbs and Grafton (2015), thus allowing a more direct comparison of repetition suppression results. Even when the error trials were excluded (i.e. removing all error trials as well as second execution trials when the first execution was erroneous), our results remained unchanged. Ultimately, the first level analysis resulted in activation images (beta maps) for each of the 24 conditions per run, for each of the four scanning sessions.

### Surface reconstruction and regions of interest

Individual subject’s cortical surfaces were reconstructed using FreeSurfer (Dale et al., 1999), and aligned to the FreeSurfer’s Left-Right symmetric template (Workbench’s 164 nodes template) via spherical registration. To examine sequence representation across the cortical surface, we defined searchlights (Oosterhof et al., 2011). A searchlight was defined for each surface node, encompassing a circular neighbourhood region containing 120 voxels. As a slightly coarser alternative to searchlights, we also defined a regular tessellation of the cortical surface separated into small hexagons.

For our regions of interest (ROI), we defined areas covering the primary motor cortex and secondary associative regions. The primary motor cortex (M1) was defined using probabilistic cytoarchitectonic map (Fischl et al., 2008) by including nodes with the highest probability of belonging to Brodmann area (BA) 4 which in addition corresponded to the hand knob area (Yousry et al., 1997). The dorsal premotor cortex (PMd) was included as the lateral part of the middle frontal gyrus. The anterior part of the superior parietal lobule (SPLa) was defined to include anterior, medial and ventral intraparietal sulcus.

### Evoked activation and repetition suppression

We calculated the percent signal change for execution of each sequence relative to the baseline activation for each voxel. The calculation was split between the first and second execution (**Fig 1d**).

To calculate repetition suppression, the activation during the first execution was subtracted from the elicited activation during the second execution. Thus, negative values of this difference contrast represented relative suppression of activation on the second execution, i.e. repetition suppression. For most subsequent analyses, the obtained values of activation and repetition suppression were averaged separately for trained and the untrained sequences. For ROI analysis, the volume maps were averaged across the predefined regions (M1, PMd, SPLa) in the native volume space of each subject. Additionally, for visualization the volume maps were projected to the surface for each subject, and averaged across the group in Workbench space.

### Dissimilarities between activity patterns for different sequences

To evaluate which regions displayed sequence-specific representation, we calculated Crossnobis dissimilarities between the evoked beta patterns of individual sequences. To do so, we first multivariately prewhitened the beta values – i.e. we standardized them by voxels’ residuals and weighted by the voxel noise covariance matrix. We used optimal shrinkage towards a diagonal noise matrix following the Ledoit and Wolf (2004) procedure. Such regularized prewhitening has been found to increase the reliability of dissimilarity estimates (Walther et al., 2016). Next, we calculated the crossvalidated Mahalanobis dissimilarities (i.e. the Crossnobis dissimilarities) between evoked regional patterns of different pairs of sequences, resulting in a total of 66 dissimilarities. This was performed twice: once by combining the activation patterns across the two executions and second time by separately obtaining dissimilarities between evoked patterns split per execution. The obtained dissimilarities were then averaged overall, as well as separately within the pairs of trained sequences, and the untrained sequences. This analysis was conducted separately for each ROI and using a surface searchlight approach (Oosterhof et al., 2011). In the searchlight approach, dissimilarities were calculated amongst the voxels of each searchlight, with the resulting dissimilarities values assigned to the centre of the searchlight.

### Changes in dissimilarities with repetition

We then related the change in dissimilarities with repetition to the changes in overall activity. As a starting point, we considered the possibility that repetition suppression simply scaled the entire activity pattern downward. To test for this possibility, we first computed the ratio of activation change: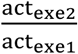. Based on this value, we could compute what dissimilarities would be predicted on the second execution if representation decreased proportional to the decrease in activation 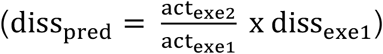. This was then contrasted with the observed dissimilarities on execution 2 (diss_exe2_ − diss_pred_). A positive difference indicates that dissimilarities decrease relatively less with repetition than the reduction in average activation. This would indicate a relatively sharper representation on the second execution. In contrast, a negative difference would reflect a further reduction in dissimilarities relative to that obtained in activation. This would suggest that with repetition, representation decreases relatively more than activation.

### Pattern component analyses: modelling representational components

To determine what specific features of the patterns might change across the two executions, we decomposed the pattern component modelling toolbox (PCM; Diedrichsen et al., 2011, 2017). PCM models the covariance structure (second moment matrix) of regional activity patterns according to different representational hypotheses.

In our experiment based on presented sequences, we defined five representational components.

#### 1) First finger

Both trained and untrained sequences started with one of three possible fingers: thumb, middle or little finger. The first finger component predicts that activity pattern for sequences that start with the same finger are identical. For sequences starting with a different first finger, the prediction was based on the covariance of the natural statistics of hand movement (Ejaz et al., 2015).

#### 2) All fingers

The sequences were slightly different in terms of which fingers were involved. The ‘all fingers’ component simply characterized how often each finger occurred in each sequence. If two sequences consisted exactly of the same presses (just in a different order), they were predicted to be identical. The predicted covariance was again weighted by the natural statistics of hand movement (Ejaz et al., 2015).

#### 3) Sequence type

This component split the performed sequences based on whether they were trained or untrained, predicting one regional activity patterns for all the trained and a different activity pattern for all the untrained sequences.

#### 4) Trained sequence identity

This component modelled any differences between the 6 trained sequences.

#### 5) Untrained sequence identity

Similar as the trained sequence identity, this component predicted a unique activity patterns for each untrained sequence.

The overall predicted second moment matrix (G) was then a convex combination of the component matrices (G_c_), each weighted by a positive component weight exp (Θ_i_).

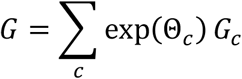

The construction of the model components was done separately for the two groups of participants, as different sequences constituted ‘trained’ or ‘untrained’ sequences for the two groups. The subsequent steps of model fitting and evaluation were carried together for all subjects.

We formulated a model family containing all possible combinations of the five chosen components (Yokoi and Diedrichsen, 2019). This resulted in 32 combinations, also containing the ‘null’ model that predicted no differences amongst any of the sequence patterns. We evaluated all of the 32 models using a crossvalidated leave-one-subject-out scheme. The components weights were fitted to maximize the likelihood of the data the data of subject 1,…,N-1. We then evaluated the likelihood of the observed regional activity patterns of subject N under that model. The resultant cross-validated likelihoods were used as model evidence for each model (see Diedrichsen et al. 2017). The log model Bayes Factor BF_m_, the difference between the crossvalidated log-likelihood of each model and the null model, characterises the relative evidence for that model.

In addition to the model family of the chosen components, we also fit a ‘noise-ceiling’ model to assess maximal logBF_m_ that would be achievable for a group model (Nili et al., 2014; Diedrichsen et al., 2017). For each of the two groups, we predicted the second moment matrix of a left-out subject based on n-1 subjects in the same group. This metric of inter-subject consistency was then combined across the subjects of the two groups.

To integrate the results across models, we used model averaging. Assuming a uniform prior probability across models, we first computed the posterior probability of each model and region directly from the log-Bayes factors:

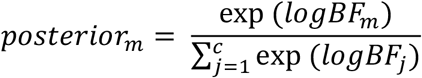

The posterior probability was used to calculate two subsequent metrics: 1) component log-Bayes factor, and 2) variance accounted for by each component. The log-Bayes factor for each component (first finger, all fingers, etc.) was calculated as the log of the ratio between the posterior probability for the models containing the component (c=1) versus the models that did not (c=0).

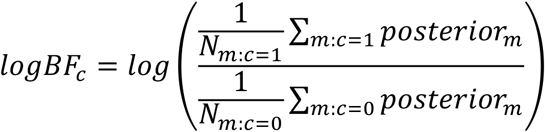

where N_m:c=1_ (N_m:c=0_) denotes the number of models (not) containing the component (Shen and Ma, 2019). The component log-Bayes factor is monotonically related to the posterior probability of model components.

To determine the amount of pattern variance accounted for by each component (across the models), we normalized the trace of each model component to be 12 (number of conditions) prior to fitting. Thus, the fitted component weight exp (Θ_i,m_) indicates the amount of variance accounted for by the component *i* in the context of model *m*. The model-averaged amount of variance accounted for by each component c was then calculated as:

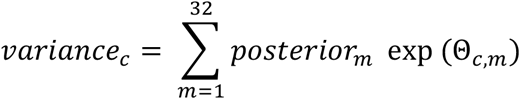

Important to note is that the estimated variance is always positive, such that this quantity cannot be used to test whether a component is present at all. On the other hand, the log-Bayes factor does not take into account the actual weighting of the component in explaining the activity patterns. In univariate models, the average variance accounted for is tightly related to the evidence for that component-however this is not necessarily the case in the multivariate setting. While component *c1* can be crucial to account for the covariance between the patterns, it may actually play a relative small role in predicting the activity patterns. Thus, both the component Bayes factor and the averaged explained variance provide informative, albeit slightly different, measures of the importance of a component.

### Statistical analysis of repetition suppression and dissimilarities

We employed a within-subject design. For each subject’s data, we calculated repetition suppression (RS) and dissimilarities, separately for trained and untrained sequences. This was done for each region and session. To statistically quantify how RS and dissimilarities changed with learning (across sessions for trained / untrained sequences), we performed a session x sequence type ANOVA on those metrics, in predefined ROIs. Afterwards, we used a two-sided paired t-test to assess the effect of sequence type per session. We additionally performed a three-way session x region x sequence type ANOVA to examine if the learning-related effects differed across regions. For the analysis of dissimilarities split by execution (execution 1 vs. 2), we calculated, per subject, the expected crossnobis dissimilarities for execution 2 of the cortical surface regions. The observed dissimilarities on the second execution were contrasted with those by using a two-sided paired t-test.

### Statistical analysis of pattern component modelling

We report the component log-Bayes factors, averaged across subjects. Additionally, the log-Bayes factors were submitted to a one-sample t-test against 0 (two-sided). To quantify the change in component variance across executions, we calculated, per subject, the percent reduction in component variance from execution 1 to 2. The relative reduction in variance with repetition was contrasted across components by using a two-sided paired t-test.

## Results

### Changes in repetition suppression with learning

To examine learning-related changes in repetition suppression and pattern analysis, we calculated both metrics on fMRI activation patterns both pre- and post-learning (i.e. weeks 1 and 5). Relative to rest, sequence execution activated primary motor cortex (M1), primary somatosensory cortex (S1), dorsal and ventral premotor cortex (PMd and PMv), supplementary motor area (SMA) and the anterior superior parietal lobules (SPLa; **Fig 2a**). In general, activity was higher for the first than for the second execution (**Fig 2b**). Repetition suppression was calculated as the difference in activity between the two executions of the same sequence (Exe 2 – Exe 1). Negative values indicate a relative reduction in activation with repetition, i.e., repetition suppression (RS). Already in week 1, prior to learning, RS was observed in nearly all regions displaying task-evoked activation (**Fig 2c**). Only in regions that showed de-activation during task performance (blue shades in **Fig 2b**), did we observed positive difference values between the executions (areas in red shades in **Fig 2c**). This indicates that, both the amount of activation and the amount of deactivation reduced with repetition.

**Figure 2.**
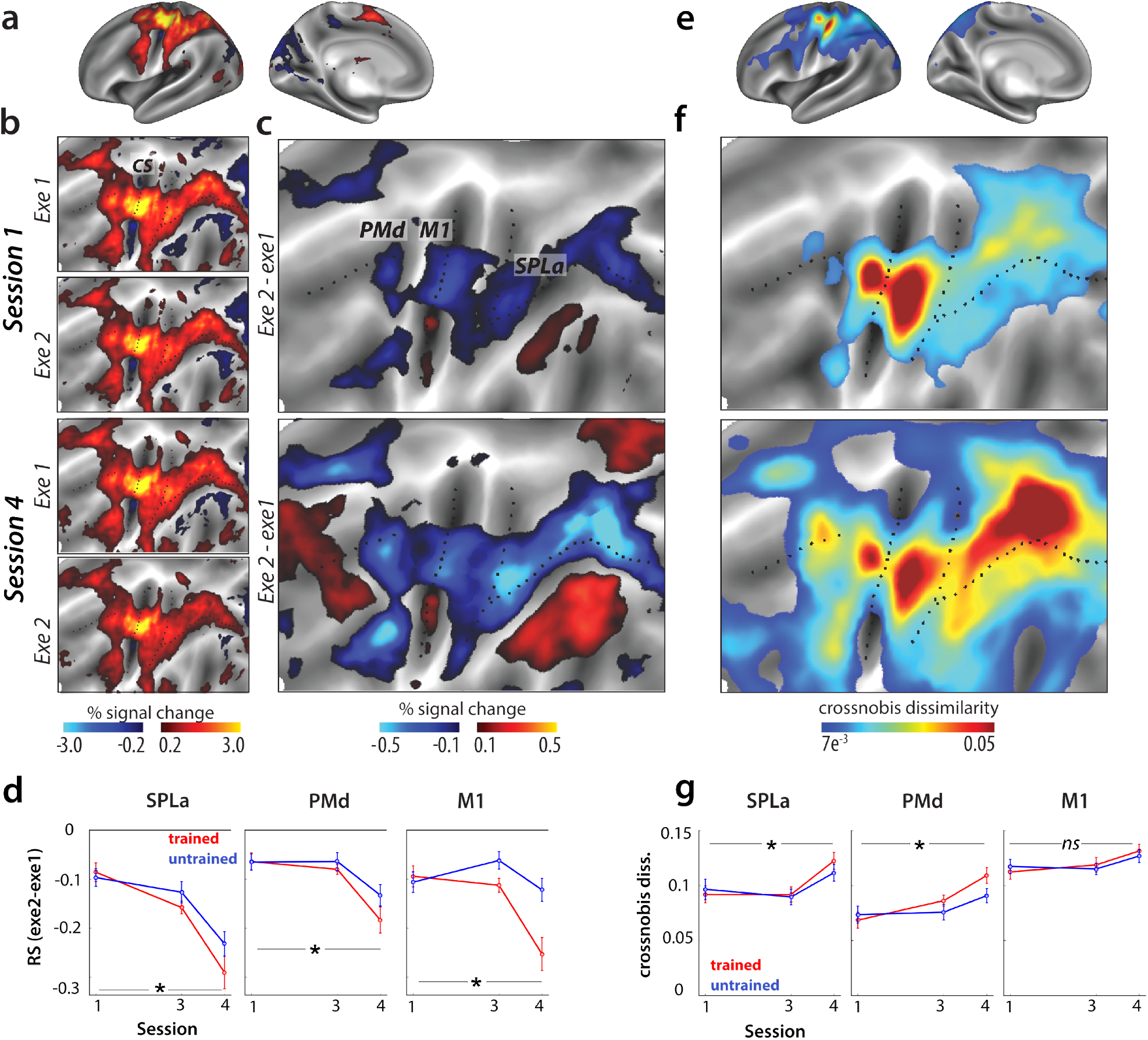
Changes in repetition suppression and dissimilarities with learning. **a)** Group-averaged evoked activation, measured as percent signal change over resting baseline in week 1, averaged across all sequences and projected to an inflated representation of the left hemisphere, i.e. hemisphere contralateral to the performing hand. **b)** Group-averaged activation for each execution (Exe1, Exe2), in the baseline session (Session 1 – Week 1) and after training (Session 4 – Week 5) represented on a flattened representation of the left hemisphere. CS stands for the central sulcus. **c)** The difference in evoked activation between the two executions. Blue represents relative suppression of activation on the second, relative to the first, execution. Regions of interest: primary motor cortex (M1), dorsal premotor cortex (PMd), anterior superior parietal lobule (SPLa). **d)** Repetition suppression in the predefined regions of interest, separately for trained (red) and untrained (blue) sequences. Error bars reflect the standard error of the group. More negative values indicate more suppression during second execution, relative to the first. * signals *p*<.05. **e)** Average dissimilarity between evoked patterns for all pairs of sequences, in week 1, averaged across the group. Pattern dissimilarity was computed using a searchlight approach, by calculating the average crossnobis dissimilarity of activation patterns between all sequence pairs in each searchlight. **f)** Average dissimilarity between activation patterns of different sequence pairs in weeks 1 and 4. **g)** Dissimilarities between trained (red) and untrained (blue) sequence patterns, across weeks 1 and 5. Error bars reflect the standard error of the group. * signals *p*<.05.

We statistically quantified how RS changed across weeks (specifically between sessions 1 and 4) for three predefined regions of interest: SPLa, PMd, and M1. The increase in RS across session was higher for trained than untrained sequences in all regions (**Fig 2d**), as confirmed by significant session x sequence type interactions (SPLa: *F*_(1,25)_=17.44; *p*=3.1e^−4^, PMd: *F*_(1,25)_=7.27, *p*=1.1e^−6^, *F*_(1,25)_=25.09; *p*=3.6e^−4^). The increase in RS was particularly strong in M1. Indeed, the three-way interaction of region x session x sequence type was significant (*F*_(2,50)_=9.19, *p*=3.9e^−4^). To summarize the RS results, all regions showed evidence of an increase of sequence-specific representation with learning, with a particularly strong effect in M1.

### Changes in pattern dissimilarities with learning

As another measure of sequence-specific representations, we tested whether the regions that displayed RS also showed distinguishable fine-grained activity patterns for each sequence. As a measure of pattern dissimilarity, we calculated the average crossvalidated Mahalanobis dissimilarity (i.e., crossnobis dissimilarity) between activation patterns of all possible sequence pairs. Overall, regions with dissimilar activity patterns for the different sequences corresponded to regions which also exhibited RS effects (**Fig 2e-f**). Additionally, both metrics (RS and pattern dissimilarities) increased from session 1 to 4, with the effect particularly pronounced in the parietal cortex (**Fig 2c, f**). Thus, based on visual inspection, RS and pattern dissimilarity metrics seem to provide consistent evidence for the development of sequence-specific representations with learning in an overlapping set of regions. However, when quantifying the change in pattern dissimilarities across weeks in predefined ROIs, we observed important differences from RS. In SPLa and PMd, pattern dissimilarities increased more for trained than untrained sequences across sessions (**Fig 2g**), as quantified by a significant interaction in a session x sequence type ANOVA (SPLa: *F*_(1,25)_=4.80; *p*=.038, PMd: *F*_(1,25)_=5.29, *p*=.030). In contrast, the week by sequence type interaction was not significant in M1 (**Fig 2g**; *F*_(1,25)_=2.13, *p*=.16). This indicates that while PMd and SPLa show learning-related changes on the level of pattern dissimilarities, these are absent in M1. The three-way interaction (region x session x sequence type) on the observed dissimilarities was indeed significant (*F*_(2,50)_=3.39, *p*=0.041), confirming the difference between regions.

### Pattern dissimilarities reduce with repetition

Within the same dataset, we observed learning-related changes in RS in M1, but no change in pattern dissimilarities with learning. While the increase in pattern dissimilarities (**Fig 2f**), as well as direct evidence for pattern changes across weeks (Berlot et al., 2020), clearly argue that sequence-specific learning occurs in premotor and parietal areas and not in M1, RS provides evidence for the development of sequence-specific representations in all these regions. How can this discrepancy be explained? To resolve this question, we need to understand how the role that each area plays during skilled sequence performance changes from the first to the second execution. We first inspected pattern dissimilarities for each of the two executions separately (execution 1, execution 2) in the trained state (Week 5 / Session 4). We observed that, on average, pattern dissimilarities in week 5 decreased with repetition in most cortical regions (**Fig 3a**). This decrease was particularly pronounced around the central sulcus, including M1 (**Fig 3b**).

**Figure 3.**
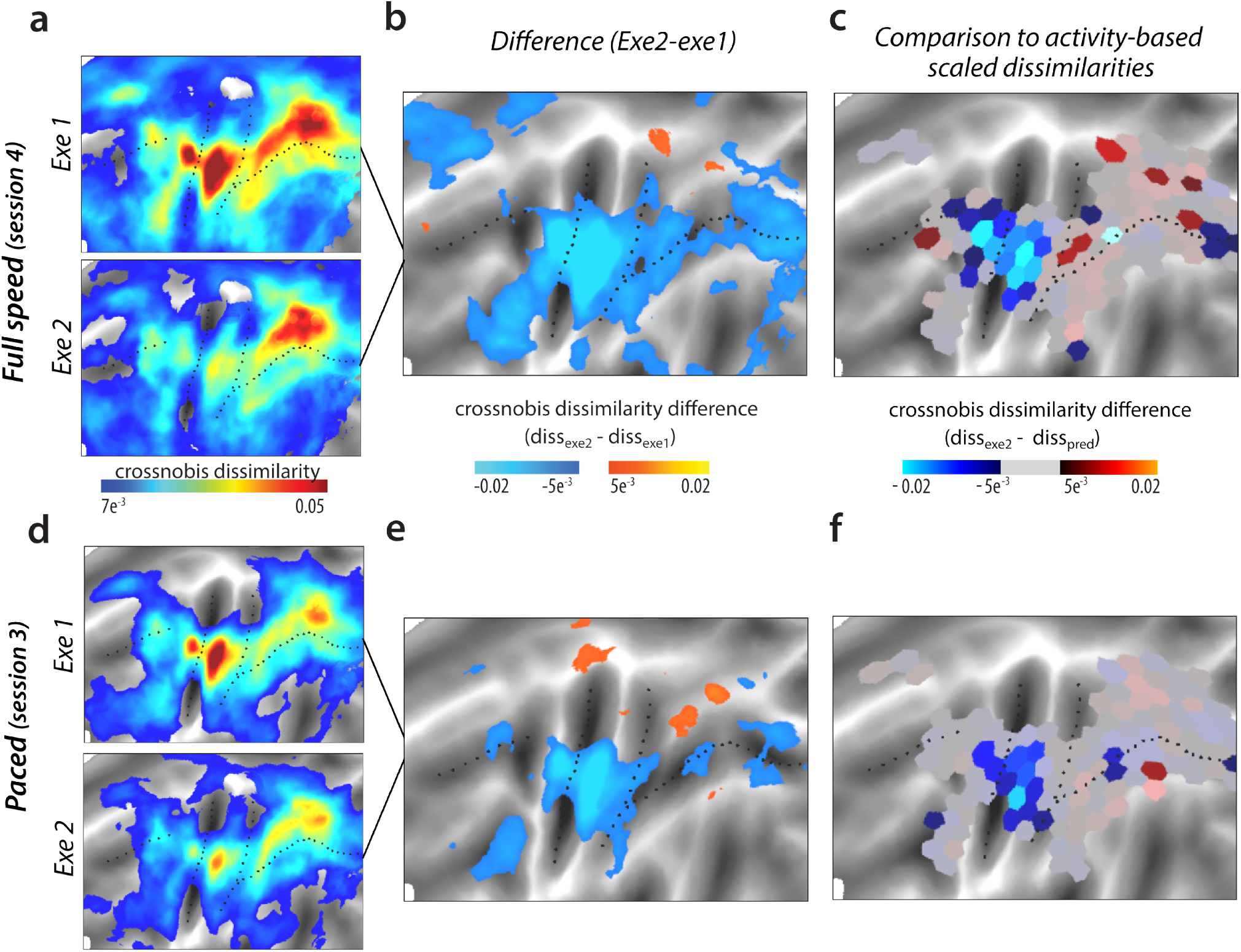
Representational change with repetition of sequence execution. **a)** Dissimilarities between pairs of sequences in session 4, split by first and second executions. **b)** Difference in pattern dissimilarities between executions 1 and 2. Blue hues reflect relatively lower dissimilarities on the second execution. **c)** Difference between the observed dissimilarity during execution 2 and the predicted distance based on the reduction of activation with repetition. Blue hues indicate lower dissimilarities than predicted, red higher. The difference between the two was significant with *p*<.05 in tessels which are fully visible (i.e. not greyed out). **d-f)**: Same as **a-c** but for the paced speed session, i.e. session 3. Same thresholds were applied to the visualizations as the respective figures from a-c.

**Figure 4.**
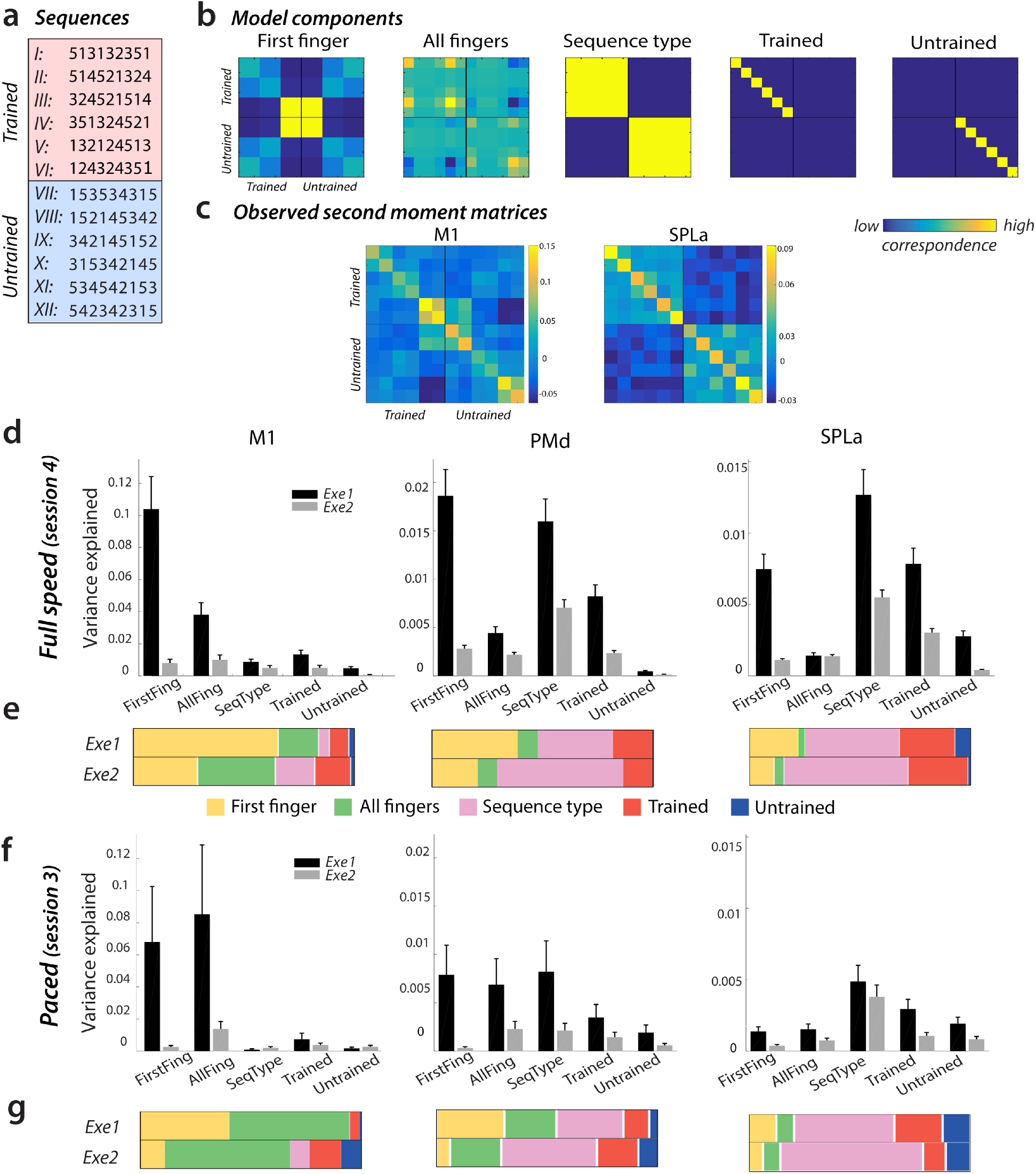
Component decomposition of regional representation across executions 1 and 2. **a)** Executed 9-digit sequences. **b)** Candidate component models used to assess regional representations across first and second executions. Each row and column indicate a specific sequence, and values in the matrices reflect the correspondence across sequences on that component, with yellow indicating higher correspondence. **c)** Regional representations during the first execution of sequences, as assessed by the crossvalidated second moment matrix, averaged across subjects of group 1. Similar as for models, each row and column reflect an activation pattern for an individual sequence. Regions: primary motor cortex (M1) and anterior superior parietal lobule (SPLa). **d)** Variance explained by candidate model components on executions 1 (black) and 2 (grey) during the full speed session in M1, PMd (dorsal premotor cortex) and SPLa. **e)** Relative contribution of variance explained in d) across the different components. The total variance explained across the different components (i.e. sum of the bars in d) was normalized across the two executions to display the relative shift of importance of different representational components. **f-g)**: Same depiction as **d-e** for the results of activity patterns during the paced scanning session.

Of course some decrease in dissimilarities would be expected given the decrease of overall activity with repetition (**Fig 2d**). We therefore compared the decrease in dissimilarities to what would be predicted if activation decreased proportionally for all sequences. First we calculated the relative decrease in activity – i.e. the ratio of the activity during the second execution over the activity during the first. This ratio was applied to the observed dissimilarities on the first execution, yielding a prediction of what dissimilarities would be expected for the second execution, if representation scaled with activation. This calculation was applied to activity patterns to each of the parcels on a regularly tessellated cortical surface (**Fig 3c**). Around the central sulcus, i.e. including M1, the observed dissimilarities on the second execution were significantly lower than what was predicted from the reduction in overall activity (**Fig 3c**). In contrast, observed dissimilarities on the second execution in premotor and parietal areas were quite close to the prediction from activation reduction. Altogether this indicates that representational change with repetition differed across regions: proportional scaling of representation in parietal regions, and violation of proportional scaling in M1, where a much more pronounced decrease of dissimilarities was observed.

### Decomposing representations across executions 1 and 2

Analysis of average dissimilarities across executions revealed a compression of representation in M1, but not in parietal regions. This analysis, however, does not reveal which aspects of the representations are responsible for this regional difference. To investigate exactly how the representation changed, we decomposed the representations during each execution into several underlying representational components. Differences in the sequence patterns could reflect differences in various characteristics, or features (**Fig 4a**). Specifically, based on previous results (Yokoi et al., 2018; Yokoi and Diedrichsen, 2019), we hypothesized that the covariance (or similarity) between activity patterns can be explained with the following 5 components (**Fig 4b**, see **Methods** for details): *1) first finger:* a pattern component determined by the starting finger, *2) all fingers*: a pattern component that simply adds the finger-specific patterns regardless of their sequence, *3) sequence type*: trained and untrained sequences have different average patterns, *4) trained sequence identity*: the trained sequences differ amongst each other, *5) untrained sequence identity*: the untrained sequences differ amongst each other. Using pattern component modelling (Diedrichsen et al., 2017), we constructed a model family, which consisted of all possible combinations of those 5 components, totalling 2^5^ = 32 models. These models were then fit to the observed regional covariance structure (second moment matrices; **Fig 4c**), separately for executions 1 and 2. In all regions and across both executions, several models accounted for observed data well, with model fits as good as the noise ceiling model (M1: 21 models for exe 1, 24 for exe 2; PMd: 16 for exe 1 and 2, SPLa: 16 for exe 1 and 2), showing that overall these models accounted well for the observed data. To integrate the results across models, we used Bayesian model averaging to estimated which components were most important to explain the patterns. In M1, the regional representation on the first execution was accounted for by the individual movement elements (all fingers), with especially high weight on the first finger (**Fig 4d**). This replicates the previous findings showing that M1’s representation during sequence production tasks can be fully explained by the starting finger (Yokoi et al., 2018; Yokoi and Diedrichsen, 2019). In these two studies, the number of times each of the five fingers was pressed was held constant across all sequences. In the current study, we did not match this number. Thus, the subsequent finger presses, encoded in the ‘all finger’ component, also accounted for substantial variance, independent of the exact ordering of these movements.

To statistically quantify these effects, we calculated component Bayes factors for individual components. In M1, the Bayes factors were significant for both first and all finger factors (first finger: BF=6.8, *t*_(25)_=3.1, *p*=4.8e^−3^; all fingers: BF=9.6, *t*_(25)_=4.4, *p*=1.7e^−4^). In contrast, the component Bayes factors were not significant for any sequence-related feature – neither sequence type (BF_c_=3.2, *t*=1.9, *p*=.07), nor sequence identity: of trained sequences (BF_c_=1.6, *t*_(25)_=1.5, *p*=.16) or untrained sequences (BF_c_=0, *t*_(25)_=-0.2, *p*=.85). Thus, the pattern analysis clearly shows that activity patterns during the first execution in M1 can be explained by a superposition of individual movements, without any evidence of a sequence representation.

In SPLa and PMd, the variance explained during the first execution was well accounted for by sequence type (SPLa: BF_c_=16.3, *t*_(25)_=6.0, *p*=3.0e^−6^, PMd: BF=15.5, *t*_(25)_=5.94, *p*=3.3e^−4^), and trained sequence identity (SPLa: BF_c_=5.4, *t*_(25)_=3.4, *p*=2.5e^−3^; PMd: BF_c_=4.6, *t*_(25)_=2.8, *p*=.011). There was no significant evidence for representation of untrained sequence identity in either of the regions (SPLa: BF_c_=0.8, PMd: BF=0.1; *t*_(25)_<=1.1, *p*>=.28). In comparison to M1, the variance related on individual movements either the first finger or all fingers were weaker across PMd and M1. In PMd the first finger still accounted for some variance (BF_c_=4.1), but this was further reduced in SPLa (BF_c_=0.5).

In M1, the pattern component related to the first finger drastically reduced by 93% with repetition (**Fig 4d**). The reduction in variance explained by the first finger component was larger than for the all finger component, which reduced by 75% (paired t-test: *t*_(25)_=9.03, *p*=2.4e^−9^). This indicates that the drastic reduction of average dissimilarities in M1 with repetition is mostly due a pronounced first-finger effect during the first execution that almost vanishes on the second execution.

Large reductions in first finger effect were also observed in session 4 in PMd (by 81%) and SPLa (by 83%). In contrast, the representation of sequence type and trained sequence identity in these areas clearly reduced less (PMd: sequence type: 44%, trained sequence: 64%; SPLa: sequence type: 49%, trained sequence: 55%). To statistically quantify whether the first finger effect reduced more than trained sequence component, we performed a paired t-tests on the percentage reduction across the two components. The results of tests were indeed significant for both PMd (*t*_(25)_=7.96, *p*=2.6^−8^) and SPLa (*t*_(25)_=12.8, *p*=1.7e^−12^).

In summary, SPLa’s regional activation patterns were better accounted for by components related to the sequence identity than to the first finger, which also reduced much less with repetition. This likely explains why the average dissimilarities did not compress with repetition in SPLa regions as much as in M1. With repetition, the proportion of different components to overall regional representation remained relatively stable in SPLa (**Fig 4e**), but changed substantially in M1 in that the dominant first-finger representation on the first execution nearly disappeared on the second execution. PMd’s representation was in-between those of M1 and SPLa – more variance was accounted for by the first finger than in SPLa, but less than in M1.

### Speed of execution does not affect RS, but it overall alters the balance between first- and all-finger representations

It is important to note that the speed of execution differed between trained and untrained sequences in session 4 (**Fig 1c**). This speed difference could conflate the observed effect of learning. To control for this factor, we had designed the study to include an extra session, session 3, which was also performed after learning was completed, but with paced performance. Specifically, the movement speed in session 3 was matched between trained and untrained sequences, as well as to performance observed in session 1.

We have previously reported that after learning, crossnobis dissimilarities for trained sequences are affected by the speed of execution. Specifically, the dissimilarities between trained sequences were lower for paced session (session 3) than full speed session 4 in PMd and SPLa, but not in M1, where there was no distinction between trained and untrained dissimilarities in either session (Berlot et al., 2020; **Fig 2g** comparison session 3-4). Similarly, RS in PMd and SPLa was also less pronounced in session 3. The RS did not differ significantly between trained and untrained sequences in session 3 (*t*_(25)_<=1.22, *p*>=.23; **Fig 2d**). However in M1, the difference in RS between the two sequence types was significant already in session 3 (*t*_(25)_=2.1, *p*=0.046). The nature of this significance is less clear since RS for neither trained nor untrained sequences changed significantly from session 1 to 3. Still, it points to the fact that the presence of learning-related effects (as characterized from session 1 to 4) in M1 for RS, but no change in dissimilarities cannot be simply explained by the speed of execution.

Next, we compared whether the speed of execution affects the decrease in dissimilarities on repetition. As for the full speed performance, we observed that dissimilarities decreased on the second execution (**Fig 3d-e**). Additionally, as reported for the full speed performance, this reduction in dissimilarities was particularly pronounced around the central sulcus (**Fig 3f**) also when performance was paced with the metronome.

Finally, we assessed whether the reduction in representational components on repetition (especially the finger effect in M1) is observed even during paced performance. Overall, our PCM modelling accounted for less variance during the paced performance compared to full speed performance (**Fig 4d,f**). We have previously reported that the patterns of activity are much more distinguishable and have higher signal-to-noise ratio during the full speed session compared to paced performance (Berlot et al., 2020), which likely accounts for this difference.

Interestingly, the overall amount of the first-vs. all-finger components varied with speed. During full speed performance the first finger component accounted for a larger part of the pattern variance than during paced performance (**Fig 4d-g**). This was confirmed by an significant interaction of a session x component (first / all fingers) ANOVA in M1 (*F*_(1,25)_=17.3, *p*=3.3e^−4^). Nevertheless, a similar reduction of the first-finger effect in M1 was observed for the paced session as for the full speed session (first finger reduction by 92%, all finger by 66%; *t*_(25)_ = 3.12, *p*=4.5e^−3^), suggesting that the decrease of the first finger weight on repetition did not depend on the speed of execution. The reductions in first finger effect were larger than for trained sequence components also in PMd and SPLa (PMd: *t*_(25)_=2.34, *p*=0.02; SPLa: *t*_(25)_=8.11, *p*=1.8e^−8^). Altogether this confirms that the larger reduction of the first finger effect with repetition does not depend on the speed of performance.

## Discussion

In the present study, we combined two fMRI analysis techniques to investigate brain underpinnings of learning motor sequences: pattern analysis and repetition suppression. Both techniques showed the development of sequence specific representations in premotor and parietal cortex. In contrast, only RS provided evidence for a sequence learning in M1. In this study, we carefully investigated how the activity patterns in these regions changed from the first to the second repetition, which offers an explanation for these discrepant findings, and which leads us to a speculative model of parietal – M1 interactions in skilled sequence performance.

### Learning-related changes of RS and pattern dissimilarities

Several pattern analysis fMRI studies have failed to provide evidence that M1 obtains a motor sequence representation with learning (Wiestler and Diedrichsen, 2013; Yokoi et al., 2018; Berlot et al., 2020). In contrast, one study (Wymbs and Grafton, 2015) reported learning-related changes in RS even for M1, which suggests a development of sequence-dependent representation. We first replicated that these two metrics provide discrepant insights into M1 – we observed evidence for learning-related changes using RS, but not pattern dissimilarities. In additional control analysis, we also showed that this difference was not due by a higher sensitivity of RS to speed of execution. The results of the session with paced performance showed that RS in M1 was stronger for trained than untrained sequences even for paced performance, whereas pattern dissimilarities did not differ between trained and untrained sequences for either full speed or paced sessions.

As Wymbs & Grafton (2015), we found changes in RS in M1 across learning sessions, as well as a difference between trained and untrained sequences in sessions post-training. However, the specific evolution of the changes differed between the two studies. Wymbs and Grafton reported a complex increase-decrease-increase pattern of RS in M1 depending on the level of the training of the sequence. In contrast, we report higher RS for trained than untrained sequences after training. There are a number of important differences in the design of the two studies which could have contributed to the observed differences in results. For instance, their design only employed full speed performance, the probability of sequence repetition was lower (50%), and the training was longer and had three groups of sequences (highly, medium, and lightly trained) rather than just two (trained and untrained). Further studies, directly manipulating any of the aforementioned differences, are needed to reconcile the findings reported here relative to the previous report of Wymbs & Grafton (2015).

### Representational changes with repetition

Reduced activity with repetition is commonly interpreted as an indication that the region represents the dimension of the stimulus along which the repetition occurred (Grill-Spector et al., 2006). For example, if a region shows less activity every time the colour of a visual stimulus repeats (rather than the shape, texture, etc.), it would provide evidence for a role of the region in the analysis of colour. However, a more complex reason for repetition suppression could be that the region’s role changes with repetition. To test for this possibility, we decomposed regional representations into different underlying components (e.g. first finger, combination of all fingers, sequence identity, etc.) separately for the first and second execution. We observed that M1 mainly represents the first finger in a sequence. This component diminishes dramatically on a repetition. In contrast, the representation of sequence type and identity, which accounted for most of the variance in parietal areas, remained more stable across the two executions. Activation patterns in PMd reflected a mixture between these two extremes. Similarly to parietal cortex, sequence type and identity components remained stable with repetition. The substantial contribution of the first finger component on the first execution, however, diminished with repetition. This suggests that PMd’s representation is a mixture of more abstract sequence representations (as in parietal regions) and representations related to single movements (as in M1). Altogether, our results suggest that RS acts differently on different components of neuronal representations. Depending on the representational composition of each region, RS can therefore be more or less pronounced.

### Interactions between cortical motor regions during sequence performance

These findings can be summarized in the following - admittedly rather speculative - model of how parietal/premotor areas and M1 interact during skilled motor sequence performance. During the first execution, premotor and parietal regions contain information about the specific sequence that needs to be executed (**Fig 5**). Premotor regions also reflect the starting finger of the sequence. These regions may send signals to M1, pre-activating the neural circuits for the movement of the first finger. This replicates a previous finding that the difference between M1’s activation patterns is explained by the starting finger, rather than true sequence representation (Yokoi et al., 2018). The finding is also consistent with results from neurophysiology (Averbeck et al., 2002) and magneto-encephalography (MEG; Kornysheva et al., 2019) showing that the first action in a sequence is most highly activated in premotor and motor areas during the preparatory period.

**Figure 5.**
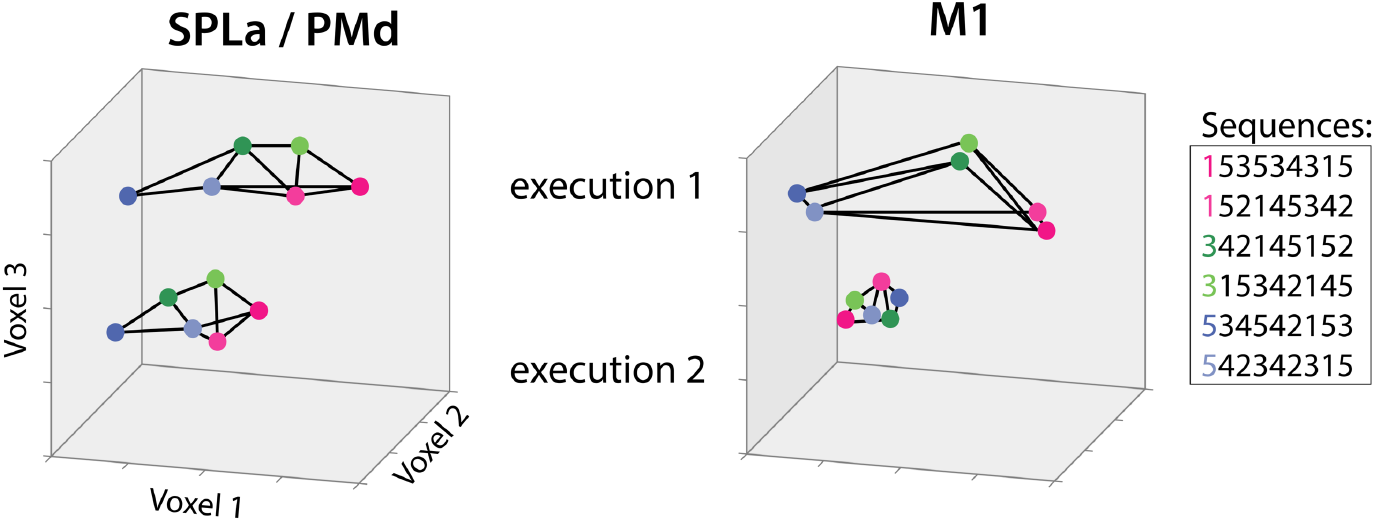
Conceptual depiction of changes in representation across regions and with repetition. Different dots represent activation patterns for different finger sequences. Regions: anterior superior parietal lobule (SPLa), dorsal premotor cortex (PMd), primary motor cortex (M1). Activation levels of three hypothetical voxels are indicated across the 3 axes.

Upon repetition of the same sequence, activation reduces across all regions. The decomposition analysis of the regional representations indicates that the sequence identity component in premotor and parietal regions reduces only moderately, suggesting that the sequence representation is always necessary for successfully guiding M1 through the correct sequences of actions. In contrast, the pre-activation of the first finger reduced dramatically, possibly reflecting reduced planning needs on repetition (Ariani et al., 2020). Thus, the especially pronounced RS effect in M1 may be due to the fact that fMRI activity here is driven to a large degree by the initial input from other regions that prepares this region for the first execution of a sequence. On the second execution, the need for this pre-activation may be substantially reduced.

Overall, our results suggest that M1 does not represent individual trained sequences with learning, despite increased RS. Instead, it appears to represent individual finger presses. If this is true, why was RS in M1 stronger for trained than untrained sequences? fMRI activity reflects a combination of the input to a cortical region, as well as the recurrent activity within that region (Logothetis, 2002), but not the output spiking (Picard et al., 2013). We suggest that the effect may be due to changed input, reflecting changes in the communication between higher-order areas and M1, which may become more efficient with repetition of trained sequences. Some support for this idea comes from a recent study demonstrating layer-specific effects in M1 (Persichetti et al., 2019). By measuring changes in cerebral blood volume across layers, the authors demonstrated that superficial M1 layers (which reflect M1 inputs) show RS, whereas deep layers’ activation (which is more indicative of M1’s outputs) is enhanced during repetition. Since the BOLD signal is biased towards the superficial vascular signals, our activation results more likely reflect inputs into M1.

However, rather than input from other areas, increased RS in M1 could reflect sequence dependency at a subvoxel resolution (Grill-Spector and Malach, 2001; Grill-Spector et al., 2006), which cannot be detected by pattern analyses. A prior electrophysiology study provided some support for this, demonstrating differential M1’s responses to trained relative to random sequences (Matsuzaka et al., 2011). However, this study did not show differential activation for different trained sequences, thus no sequence representation as defined here. Moreover, recent electrophysiological studies have also shown that M1 does not represent the sequential context (Russo et al., 2020; Zimnik and Churchland, 2021). Altogether, this makes it unlikely that the RS observed in M1 reflects sequence dependency.

Our proposed model makes a number of predictions that could be tested using a combination of techniques. For layer-specific fMRI studies, we would predict that the first finger effect in M1 can be mostly found in the superficial layers, reflecting cortico-cortico communication. For MEG or intracranial EEG studies (Ghuman et al., 2008; Gilbert et al., 2010; Korzeniewska et al., 2020) we would predict that the difference between trained and untrained sequences would be mainly present at the start of the sequence, an effect that would strongly reduce on repetition. Addressing these questions will advance our understanding of motor sequence on neural circuitry underlying production of skilled actions.

## Conclusion

We demonstrated here that RS may not only reflect a suppression of a specific representation in a region, but that the role of the region, and hence the structure of the representation, can change qualitatively from the first to the second repetition. While the representation of the skilled motor sequences remained relatively stable in parietal and premotor regions, the M1’s representation changed, with a strongly reduced activation related to the beginning of the sequence. These results emphasize that employing RS only using the average regional activation sometimes provides incomplete, and possibly misleading, insights into regional representation. Instead, the combination of RS with pattern analyses can illuminate how representations change with repetition, and may provide a deeper understanding of brain circuits and their function.

## Acknowledgements

The work was supported by an Ontario Trillium Scholarship to EB, an NSERC Discovery Grant (RGPIN-2016-04890) to JD, and the Canada First Research Excellence Fund (BrainsCAN). We thank Giacomo Ariani for helpful comments on an earlier version of the manuscript.

## References

Alink A, Abdulrahman H, Henson RN (2018) From neurons to voxels - repetition suppression is best modelled by local neural scaling. Nat Commun 9:1–10 Available at: https://www.biorxiv.org/content/early/2017/07/31/170498.

Ariani G, Pruszynski JA, Diedrichsen J (2020) Motor planning brings human primary somatosensory cortex into movement-specific preparatory states. bioRxiv:2020.12.17.423254 Available at: https://doi.org/10.1101/2020.12.17.423254 [Accessed March 24, 2021].

Averbeck BB, Chafee M V, Crowe DA, Georgopoulos AP (2002) Parallel processing of serial movements in prefrontal cortex. PNAS 99:13172–13177.

Barron HC, Vogels TP, Emir UE, Makin TR, O’Shea J, Clare S, Jbabdi S, Dolan RJ, Behrens TEJ (2016) Unmasking Latent Inhibitory Connections in Human Cortex to Reveal Dormant Cortical Memories. Neuron 90:191–203 Available at: http://dx.doi.org/10.1016/j.neuron.2016.02.031.

Berlot E, Popp NJ, Diedrichsen J (2018) In search of the engram, 2017. Curr Opin Behav Sci 20.

Berlot E, Popp NJ, Diedrichsen J (2020) A critical re-evaluation of fMRI signatures of motor sequence learning. Elife 9:1–24.

Buckner RL, Goodman J, Burock M, Rotte M, Koutstaal W, Schacter D, Rosen B, Dale AM (1998) Functional-Anatomic Correlates of Object Priming in Humans Revealed by Rapid Presentation Event-Related fMRI. 20:285–296.

Dale AM, Fischl B, Sereno MI, Dale AM (1999) Cortical Surface-Based Analysis. Neuroimage 9:179–194 Available at: http://www.ncbi.nlm.nih.gov/pubmed/9931268 http://linkinghub.elsevier.com/retrieve/pii/S1053811998903950.

Dayan E, Cohen LG (2011) Neuroplasticity subserving motor skill learning. Neuron 72:443–454 Available at: http://dx.doi.org/10.1016/j.neuron.2011.10.008.

Diedrichsen J, Ridgway GR, Friston KJ, Wiestler T (2011) Comparing the similarity and spatial structure of neural representations: A pattern-component model. Neuroimage 55:1665–1678 Available at: http://dx.doi.org/10.1016/j.neuroimage.2011.01.044.

Diedrichsen J, Yokoi A, Arbuckle SA (2017) Pattern component modeling: A flexible approach for understanding the representational structure of brain activity patterns. Neuroimage.

Ejaz N, Hamada M, Diedrichsen J ouml rn (2015) Hand use predicts the structure of representations in sensorimotor cortex. Nat Neurosci 18:1034–1040 Available at: http://dx.doi.org/10.1038/nn.4038 papers3://publication/doi/10.1038/nn.4038.

Fischl B, Rajendran N, Busa E, Augustinack J, Hinds O, Yeo BTT, Mohlberg H, Amunts K, Zilles K (2008) Cortical folding patterns and predicting cytoarchitecture. Cereb Cortex 18:1973–1980.

Friston KJ, Worsley KJ, Poline J-PB, Frith CD, Frackowiak RSJ, Holmes a. P, Worsley KJ, Poline J-PB, Frith CD, Frackowiak RSJ (1994) Statistical parametric maps in functional imaging: A general linear approach. Hum Brain Mapp 2:189–210 Available at: http://doi.wiley.com/10.1002/hbm.460020402.

Fritsche M, Lawrence SJD, de Lange FP (2020) Temporal tuning of repetition suppression across the visual cortex. J Neurophysiol 123:224–233.

Grafton ST, Hamilton AFDC (2007) Evidence for a distributed hierarchy of action representation in the brain. 26:590–616.

Grill-Spector K, Henson R, Martin A (2006) Repetition and the brain: Neural models of stimulus-specific effects. Trends Cogn Sci 10:14–23.

Grill-Spector K, Malach R (2001) fMRI-adaption: A tool for studying fucntional properties of human cortical neurons. Acta Psychol (Amst) 107:293–321.

Gross, C.G., Schiller, P.H., Wells, C., Gerstein GL (1967) Single-unit activity in temporal association cortex of the monkey. J Neurophysiol 30:833–843.

Henson RN, Ganel T, Otten LJ (2003) Electrophysiological and Haemodynamic Correlates of Face Perception, Recognition and Priming. :793–805.

Kok P, Jehee JFM, de Lange FP (2012) Less Is More: Expectation Sharpens Representations in the Primary Visual Cortex. Neuron 75:265–270.

Kornysheva K, Bush D, Meyer SS, Sadnicka A, Barnes G, Burgess N (2019) Neural Competitive Queuing of Ordinal Structure Underlies Skilled Sequential Action. Neuron 101:1166–1180.e3.

Lashley K (1950) In search of the engram.

Ledoit O, Wolf M (2004) Honey, I shrunk the sample covariance matrix. J Portf Manag 30:110–119 Available at: https://jpm.pm-research.com/content/30/4/110 [Accessed March 24, 2021].

Logothetis NK (2002) The neural basis of the blood-oxygen-level-dependent functional magnetic resonance imaging signal. Philos Trans R Soc B Biol Sci 357:1003–1037.

Matsuzaka Y, Picard N, Strick PL, Barnes TD, Mao J, Hu D, Kubota Y, Dreyer A a, Brown EN, Graybiel AM, Kilavik BE, Confais J, Ponce-alvarez A, Diesmann M, Riehle A (2011) Skill Representation in the Primary Motor Cortex After Long-Term Practice Skill Representation in the Primary Motor Cortex After Long-Term Practice. J Neurophysiol:1819–1832.

Nili H, Wingfield C, Walther A, Su L, Marslen-Wilson W, Kriegeskorte N (2014) A Toolbox for Representational Similarity Analysis. PLoS Comput Biol 10:e1003553.

Noppeney U, Price CJ (2004) An fMRI Study of Syntactic Adaptation. J Cogn Neurosci 16:702–713.

Oosterhof NN, Wiestler T, Downing PE, Diedrichsen J (2011) A comparison of volume-based and surface-based multi-voxel pattern analysis. Neuroimage 56:593–600 Available at: http://dx.doi.org/10.1016/j.neuroimage.2010.04.270.

Persichetti AS, Avery JA, Huber L, Merriam EP, Martin A (2019) Layer-Specific Contributions to Imagined and Executed Hand Movements in Human Primary Motor Cortex. SSRN Electron J:1–5.

Picard N, Matsuzaka Y, Strick PL (2013) Extended practice of a motor skill is associated with reduced metabolic activity in M1. Nat Neurosci 16:1340–1347 Available at: http://www.pubmedcentral.nih.gov/articlerender.fcgi?artid=3757119&tool=pmcentrez&rendertype=abstract.

Poldrack RA (2000) Imaging brain plasticity: Conceptual and methodological issues - A theoretical review. Neuroimage 12:1–13.

Russo AA, Khajeh R, Bittner SR, Perkins SM, Cunningham JP, Abbott LF, Churchland MM (2020) Neural Trajectories in the Supplementary Motor Area and Motor Cortex Exhibit Distinct Geometries, Compatible with Different Classes of Computation. Neuron 107:745–758.e6 Available at: https://doi.org/10.1016/j.neuron.2020.05.020.

Segaert K, Weber K, de Lange FP, Petersson KM, Hagoort P (2013) The suppression of repetition enhancement: A review of fMRI studies. Neuropsychologia 51:59–66 Available at: http://dx.doi.org/10.1016/j.neuropsychologia.2012.11.006.

Shen S, Ma WJ (2019) Variable precision in visual perception. Psychol Rev 126:89–132 Available at: /pmc/articles/PMC6318066/ [Accessed March 24, 2021].

Shmuelof L, Zohary E (2005) Dissociation between Ventral and Dorsal fMRI. 47:457–470.

Summerfield C, Trittschuh EH, Monti JM, Mesulam MM, Egner T (2008) Neural repetition suppression reflects fulfilled perceptual expectations. Nat Neurosci 11:1004–1006.

Walther A, Nili H, Ejaz N, Alink A, Kriegeskorte N, Diedrichsen J (2016) Reliability of dissimilarity measures for multi-voxel pattern analysis. Neuroimage 137:188–200 Available at: http://dx.doi.org/10.1016/j.neuroimage.2015.12.012.

Weiner KS, Sayres R, Vinberg J, Grill-Spector K (2010) fMRI-adaptation and category selectivity in human ventral temporal cortex: Regional differences across time scales. J Neurophysiol 103:3349–3365.

Wiestler T, Diedrichsen J (2013) Skill learning strengthens cortical representations of motor sequences. Elife 2013:1–20.

Wymbs NF, Grafton ST (2015) The human motor system supports sequence-specific representations over multiple training-dependent timescales. Cereb Cortex 25:4213–4225.

Yokoi A, Arbuckle SA, Diedrichsen J (2018) The role of human primary motor cortex in the production of skilled finger sequences. J Neurosci 38:1430–1442.

Yokoi A, Diedrichsen J (2019) Neural Organization of Hierarchical Motor Sequence Representations in the Human Neocortex. Neuron 103:1178–1190.e7.

Yousry TA, Schmid UD, Alkadhi H, Schmidt D, Peraud A, Buettner A, Winkler P (1997) Localization of the motor hand area to a knob on the precentral gyrus. A new landmark. Brain 120:141–157.

Zimnik, Andrew J., Churchland MM (2021) Independent generation of sequence elements by motor cortex. Nat Neurosci:1–13.

